# Simultaneous evolution of multiple dispersal components and kernel

**DOI:** 10.1101/037606

**Authors:** Sudipta Tung, Abhishek Mishra, P.M. Shreenidhi, Mohammed Aamir Sadiq, Sripad Joshi, V.R. Shree Sruti, Sutirth Dey

## Abstract

Global climate is changing rapidly and is accompanied by large-scale fragmentation and destruction of habitats. Since dispersal is the first line of defense for mobile organisms to cope with such adversities in their environment, it is important to understand the causes and consequences of evolution of dispersal. Although dispersal is a complex phenomenon involving multiple dispersal-traits like propensity (tendency to leave the natal patch) and ability (to travel long distances), the relationship between these traits is not always straight-forward, it is not clear whether these traits can evolve simultaneously or not, and how their interactions affect the overall dispersal profile. To investigate these issues, we subjected four large (*N*∼2500) outbred populations of *Drosophila melanogaster* to artificial selection for increased dispersal, in a setup that mimicked increasing habitat fragmentation over 33 generations. The propensity and ability of the selected populations were significantly greater than the non-selected controls and the difference persisted even in the absence of proximate drivers for dispersal. The dispersal kernel evolved to have significantly greater standard deviation and reduced values of skew and kurtosis, which ultimately translated into the evolution of a greater frequency of long-distance dispersers (LDDs). We also found that although sex-biased dispersal exists in *Drosophila melanogaster*, its expression can vary depending on which dispersal component is being measured and the environmental condition under which dispersal takes place. Interestingly though, there was no difference between the two sexes in terms of dispersal evolution. We discuss possible reasons for why some of our results do not agree with previous laboratory and field studies. The rapid evolution of multiple components of dispersal and the kernel, expressed even in the absence of stress, indicates that dispersal evolution cannot be ignored while investigating eco-evolutionary phenomena like speed of range expansion, disease spread, evolution of invasive species and destabilization of metapopulation dynamics.

**Data Accessibility:** Data will be deposited to dryad if accepted.

## Introduction

Climate change (reviewed in Root, et al. 2003) and various human activities (Vitousek, et al. 1997) have led to massive habitat loss and fragmentation, which in turn have affected many natural populations all over the world. These effects include, *inter alia*, loss of biodiversity, increase in extinction probability, modified species interaction patterns within a community, decrease in the average length of the trophic chains and reduced reproductive success (reviewed in Fahrig 2003). Dispersal is one of the ways by which organisms can cope with such stresses as it allows them to increase their survival probability by tracking favorable environmental conditions (Travis, et al. 2013). As a result, evolution of dispersal and its consequences have been a major focus of research in evolutionary ecology for the last few decades (reviewed in Bowler and Benton 2005, Clobert, et al. 2012, Ronce 2007). According to the classical theoretical literature, dispersal could evolve due to three primary reasons, namely, inbreeding avoidance (Charlesworth and Charlesworth 1987), reduction of kin-competition (Gandon 1999) and spatio-temporal environmental heterogeneity (McPeek and Holt 1992). However, substantial bodies of empirical and theoretical work over the last few decades suggest that the picture may not be so simple after all (reviewed in Bonte, et al. 2012).

In terms of movement, the phenomenon of dispersal is often subdivided into three stages, namely emigration from the natal habitat, inter-patch movement and immigration into the destination patch (Bowler and Benton 2005). The environment experienced during each of these stages, and the corresponding behavioral and physiological attributes needed to tackle them, can be very different. Consequently, in terms of life-history, dispersal is actually a composite trait, made up of components like propensity (i.e. the fraction of dispersers leaving the current habitat) which is primarily related to emigration, and ability (i.e. the mean distance travelled) which is primarily related to inter-patch movement. Evidently, which component(s) of dispersal evolve(s) is contingent upon the nature of the selection pressure faced by each component, the costs associated with them, how these costs interact with each other and how they are countered by the organisms (Bonte, et al. 2012). For example, in laboratory populations of *C. elegans*, dispersal propensity evolves when patch fitness is varied by externally imposed extinctions (Friedenberg 2003). However, the same trait fails to evolve when patch fitness is altered by varying resource density (Friedenberg 2003). Similarly, spatially correlated extinctions select for long distance dispersers in the spider mite (*Tetranychus urticae*) but randomly distributed local extinctions do not (Fronhofer, et al. 2014). To complicate matters further, although the various components of dispersal are related to each other, evolution of a given component does not necessarily make an organism better in terms of another component. For example, in spider mites, artificial selection can increase dispersal propensity (Yano and Takafuji 2002) but not dispersal ability (Bitume, et al. 2011). In the same organism, when selection is imposed in the form of spatially correlated extinctions, the frequency of long distance dispersers (LDDs) increases but dispersal propensity is reduced (Fronhofer, et al. 2014).

Thus, for any organism under a given ecological scenario, a complete picture of dispersal evolution is possible only when all the dispersal components are simultaneously investigated. Unfortunately, most empirical studies typically focus on the evolution of any one of the several components of dispersal (Bitume, et al. 2011, Friedenberg 2003, Keil, et al. 2001, Ogden 1970, Tien, et al. 2011, Yano and Takafuji 2002; although see Fronhofer, et al. 2014). This makes it somewhat difficult to envisage how the evolutionary responses to the individual components of dispersal ultimately come together to affect the distribution of dispersal distances of individuals, i.e. the dispersal kernel (Nathan, et al. 2012).

The kernel is one of the most frequently used descriptors of the outcome of dispersal in the ecological and the evolutionary literature (reviewed in Nathan, et al. 2012). Empirical studies suggest that kernels in natural populations are often “fat-tailed” (Clark, et al. 2005, Van Houtan, et al. 2007). This implies that many natural populations have a larger number of long-distance dispersers or LDDs, i.e. individuals who disperse far more than the mean dispersal distance of the population than what is expected by a Gaussian function. The presence of LDDs can impact several ecological phenomena like range advance (Phillips, et al. 2008), effects of habitat fragmentation (Van Houtan, et al. 2007), invasive potential (Kot, et al. 1996) and disease spread (Rappole, et al. 2006). Thus, evolution of the kernel in general, and the fraction of LDDs in particular, is a major topic of interest in the context of dispersal evolution (reviewed in Hovestadt, et al. 2012). Unfortunately, although it is easy to conceptualize a dispersal kernel, it is not experimentally simple to measure it. Moreover, the observed dispersal kernel is a product of the phenotype of the organism and the environment through which dispersal is happening. Differentiating between these two effects is not always a straightforward task. Not surprisingly, therefore, although theoretically well-investigated (e.g. Phillips, et al. 2008, Starrfelt and Kokko 2010), we are aware of only one empirical study that has demonstrated kernel evolution (Fronhofer, et al. 2014).

Another factor that can play a role in the evolution of dispersal in a population is sex. Sex-biased dispersal (SBD) is well-documented in the animal kingdom, particularly among birds and mammals (reviewed in Pusey 1987), and also insects (Bennett, et al. 2013, Lagisz, et al. 2010). Although SBD is hypothesized to be primarily driven by the mating system of the species (Greenwood 1980), recent studies have challenged this claim (Mabry, et al. 2013, Trochet, et al. 2016). One major complication here is that SBD has often been investigated in the context of any one component of dispersal like propensity or ability (Clutton-Brock and Lukas 2012). However, the presence or absence of SBD for one dispersal component (say propensity) does not allow us to make any predictions about it in the context of another dispersal component. Moreover, potentially, the realization of SBD for a given dispersal component itself can be environment-dependent. To take a hypothetical example, one sex (for example, males) might show greater dispersal ability only under resource limitation and not when resources are plentiful. Clearly, this context-dependence of SBD has consequences for gene flow across the habitats and how that affects the standing variation (Prout 1981). It can also potentially alter the evolutionary outcome depending on whether resources were available or not during the selection process. Thus, in order to obtain a more detailed picture of SBD in a given species, it is again critical to simultaneously investigate multiple dispersal components in different environments.

To address some of the issues discussed above, we subjected four large, replicate populations of *Drosophila melanogaster* to directional selection for increased dispersal. Our selection protocol mimicked increasing habitat-fragmentation over generations, with starvation and desiccation stress being the primary inducer of dispersal. The selected populations rapidly evolved to have significantly greater dispersal propensity and ability, irrespective of the presence or absence of starvation/desiccation stress. To the best of our knowledge, this is the first report of condition-dependent selection leading to the evolution of phenotype-dependent dispersal. We then describe the impact of these evolutionary changes on the shape of the dispersal kernel and how that affected the spatial extent. We also investigate whether there was sex-bias in different dispersal components and whether selection caused the two sexes to respond differently. Finally, we briefly discuss some of the eco-evolutionary implications of our results and why some of them do not match with previous observations in the literature.

## Methods

### Ancestry of the populations

The experimental populations used in this study were derived from four independent large (breeding population size of ∼2400 adults) laboratory populations of *Drosophila melanogaster* (DB_1-4_) (see Online Appendix S1: Text S1 for details). From each DB_*i*_ population (where *i*∈[1, 4]), we derived two populations: VB_*i*_ (short for ‘**V**aga**b**ond’, subjected to selection for dispersal) and VBC_*i*_ (‘**VB**-**C**ontrol’, which experiences no selection for dispersal). Thus, VB and VBC populations that share a numerical subscript (e.g. say VB_1_ and VBC_1_) were related by ancestry (DB_1_ in this case) and hence were always assayed together and treated as blocks in statistical analyses.

### Maintenance regime of experimental populations

The detailed maintenance regimes of the experimental populations are given as Online Supplemental Experimental Procedures (Online Appendix S1: Text S1). Briefly, both VB and VBC populations underwent a 15-day discrete generation cycle at a temperature of 25°C with a breeding population size of ∼2400 adults.

### Selection for dispersal

We present a brief description of the selection set-up here with the details being relegated to the Supplemental Experimental Procedures (Online Appendix S1: Text S1 and see Fig. S1). The set-up consisted of three components named source, path and destination. The source and the destination were cylindrical, transparent plastic containers (volume ∼1.5 L) that were connected by a clear plastic tube of ∼1 cm internal diameter. 12 days post egg-collection, we introduced the adult flies into the empty source. The absence of food or moisture in the source was a potential stress as the ancestors of these flies have a mean survivorship time under desiccation of only ∼19 hrs (S. Tung personal observation). Under these conditions, a subset of the flies in the source would disperse to the destination through the path. The flies were allowed to disperse for 6 hours or until roughly 50% of the population reached the destination (whichever happened earlier). Only the flies that reached the destination were allowed to breed for the next generation and the ones who remained in the source container or path were discarded. Since the imposed selection allowed ∼50% of the flies of a population to breed, we kept the breeding population size to ∼2400 by subjecting ∼4800 flies per VB_*i*_ population to selection for dispersal. For this, we used two independent “source-path-destination” setups for each VB_*i*_ population, with ∼2400 flies in each source. Post-dispersal, the dispersed flies in the two destination containers for a given VB_*i*_ population were mixed and transferred to a population cage. The control populations (VBCs) were also introduced into the cages at the same time after being maintained identically as the VBs except for dispersal.

The length of the path for the VB populations was 2 m at the beginning of the selection but was increased intermittently over the generations. By the end of the 33^rd^ generation (when the last set of assays were performed), the length of the path had reached 10 m. Preliminary studies (unpublished data) showed that there exists notable variation with respect to the dispersal ability of the individuals in these fly populations i.e. all flies do not reach the destination even if allowed to move for a relatively longer period. Thus, the selection pressure experienced by the flies due to increasing path-length was analogous to increasing distance between the patches due to habitat fragmentation.

### Assays

All assays were performed after growing both VB and VBC populations for one generation under common conditions, to minimize the contributions of non-genetic parental effects.

### Dispersal kernel assay

This assay was used to investigate the differences in dispersal propensity and ability between the VBs and the VBCs. We performed this assay thrice: after 10, 20 and 33 generations of selection. The assay-setup was similar to the selection setup described above except for the path-length. For the 10^th^ generation assay, we had a path length of 10 m, while for the others the path length was 20 m. For all cases, the entire path was divided into several detachable sections such that the number of flies in each section (or ‘distance bin’) can be counted separately (See Online Appendix S1: Text S1). On the 12^th^ day after egg collection, ∼2000 adult flies were introduced into the source container and were allowed to disperse for six hours. The duration of six hours was chosen due to two reasons. First, this is the maximum duration allowed for dispersal during the selection process and thus our assay conditions are the same as the selection conditions. Second, during the first six hours of dispersal, mortality due to desiccation is negligible, which ensures that the measured dispersal distances are not confounded by the desiccation resistance of the flies. The source container was empty (for the 10^th^ and 20^th^ generation assay) or had ∼20 ml banana-jaggery medium (for the 33^rd^ generation assay). After dispersal, each section of the path was independently sealed with all the flies in it. The flies were then heat killed and the number of male and female flies in each section was recorded. For each VB_*i*_ and VBC_*i*_ population, there were three such replicate kernel setups. For each, dispersal propensity and ability were estimated as follows:

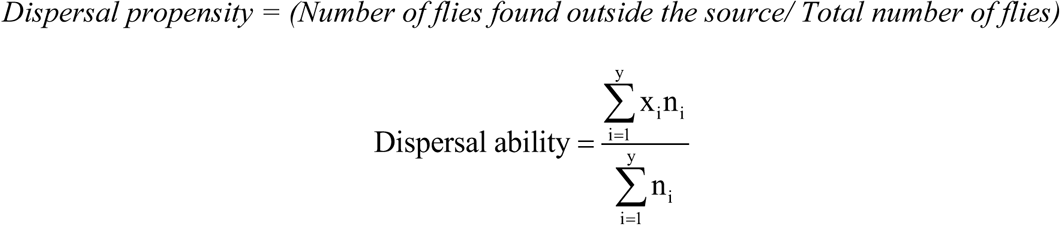

where n_i_ is the number of flies found in the i^th^ ‘distance bin’, x_i_ is the distance of the mid-point of this bin from source and y is the total number of bins. Note that we estimated dispersal ability only on the individuals that emigrated from the source. This allowed us to avoid a potentially confounding effect of lower propensity (in the VBCs) on the estimation of dispersal ability. This is because the fraction of individuals who stayed in the source (and thus had a dispersal distance of zero) was much greater for the VBCs than the VBs (Fig. 1a, 1c and 1e). If we had included the individuals in the source in our estimates, then the difference between the dispersal abilities of VBs and VBCs would have been even greater than what is represented in Figs 1b, 1d and 1f. Thus, our interpretation of increase in the dispersal ability of VBs is conservative. In total, this set of assays involved sexing and scoring of ∼140, 000 flies.

**Figure 1.**
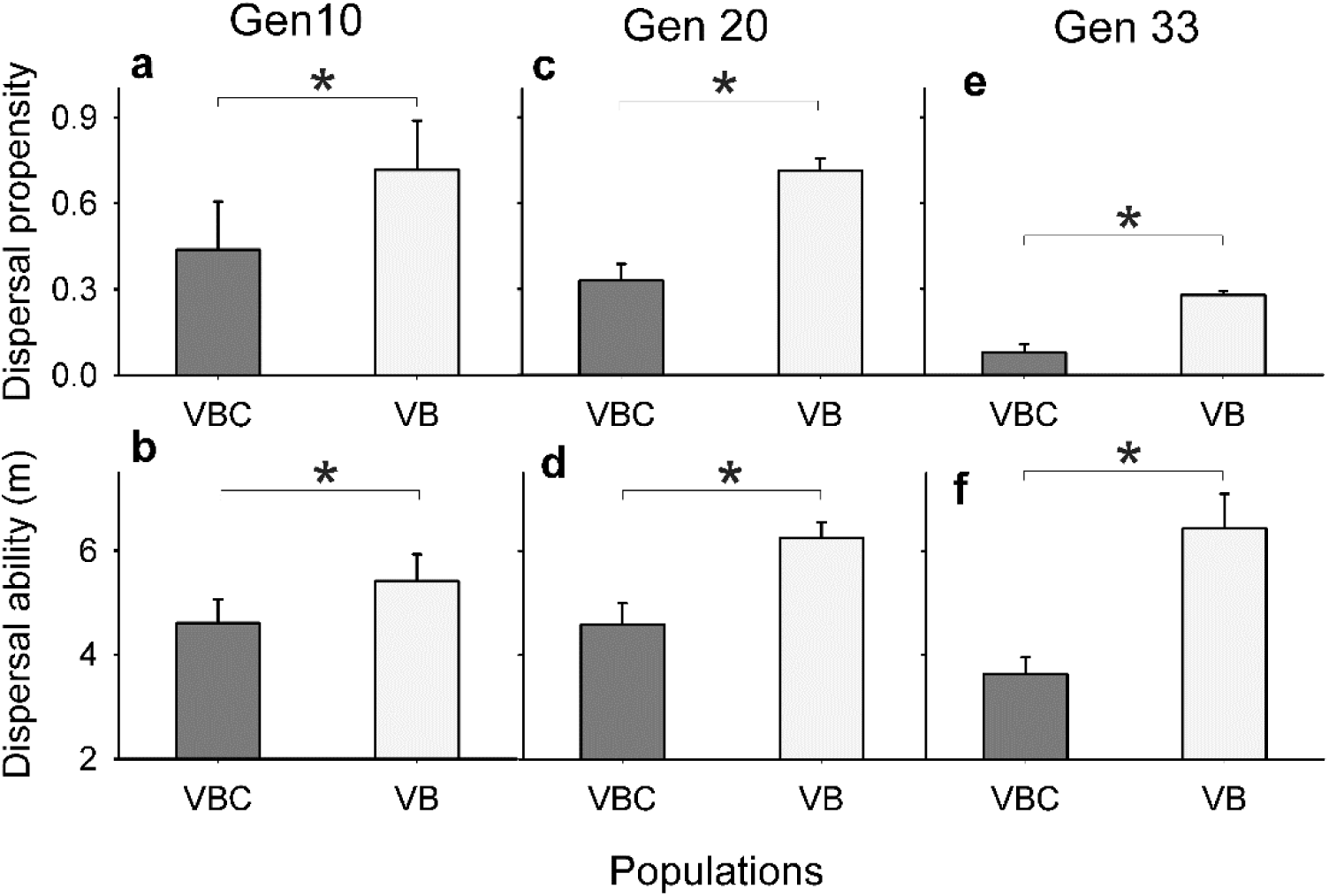
Evolution of dispersal propensity and ability. (**a, c and e**) Propensity refers to the fraction of the total population that disperses from the source. (**b, d and f**) Ability refers to the mean distance travelled by those individuals that come out of the source. The selected populations (VBs) had significantly greater propensity and ability compared to the controls (VBCs) in all the assays performed after 10 (**a, b**), 20 (**c, d**) and 33 (**e, f**) generations of selection. Food was present in the source container for the assay performed after 33 generations of selection (**e, f**). Each bar is a mean over four replicate populations each of which had three independent replicates. Error bars represent standard errors around the mean (SEM). * denotes *P*<0.05 for the main effect of selection in the ANOVA.

### Measures of dispersal kernel and spatial extent

Dispersal kernels of the VB/VBCs were characterized using the various percentiles of the distribution. Change in the mean distance travelled, in principle, can shift the kernel without changing its shape. We eliminated this effect by computing the percentiles after subtracting the mean distance travelled in a given kernel replicate from the distance travelled by each individual in the replicate. Thus for each replicate, mean-subtracted distance travelled by each individual (i.e. *x*_*i*_ − ∑*x*_*i*_/*n*), where *x_i_* is the distance travelled by *i*^th^ individual and *n* is the number of individuals that initiated dispersal in that replicate, was used for computing percentiles. To investigate shape, we calculated the higher moments of the dispersal kernel like standard deviation, skew and kurtosis (where the kurtosis of a normal distribution was taken to be zero).

To further characterize the kernel, we fit the data with the negative exponential distribution *y* = *ae^−bx^* (Kot, et al. 1996), where *x* is the distance from the source, *y* is the frequency of individuals found at *x*, and *a, b* are the intercept and slope parameters respectively. The value of spatial extent was estimated as *b*^−1^. ln (*a*/0.01), i.e. the distance from the source beyond which 1% of the population is expected to disperse. It should be noted here that in the literature, the dispersal kernel has been modeled using a large number of distributions (Nathan, et al. 2012). Thus, in principle, we could have used several other functions for fitting our kernel data. However, there were multiple reasons for which we decided to use the negative exponential distribution in this study. First, our experimental conditions were extremely well-controlled that avoided confounding factors like spatial heterogeneity or multiple dispersal agents (e.g. wind). This justifies the use of a simple kernel as opposed the large number of more complex “mixed” kernels studied in the literature (Nathan, et al. 2012). Second, the main purpose of this part of the study was to obtain an estimate of the spatial extent and the negative-exponential function performs that job well (Kot, et al. 1996). Finally, we obtained excellent fits to the data using this model (see Results and Online Appendix Table S1).

### Statistical analyses

Since VB_*i*_ and VBC_*j*_ that shared a subscript (i.e. *i* = *j*) were related to each other by ancestry, they were analyzed together as a block. Data for dispersal propensity and dispersal distance were subjected to separate three-factor mixed-model ANOVA with selection (VB and VBC) and sex (male and female) as fixed factors and block (1-4) as a random factor. The propensity data, being fractions, were arcsine-square root transformed (Zar 1999) before analysis. The standard deviation, skew, kurtosis, *a*, *b* and spatial extent data for each population were computed after pooling the data for the corresponding three measurement replicates. For these six quantities, we used separate Mann-Whitney U (MWU) tests to compare the VBs and the VBCs. The effect sizes (Cohen’s *d*) for the differences between VBs and VBCs for these six quantities were estimated. Following standard recommendations (REF Cohen 1988), the value of effect size (*d*) was interpreted as large, medium and small when d>0.8, 0.8>d>0.5 and d<0.5 respectively. All statistical analyses were performed using STATISTICA^®^ v5 (StatSoft. Inc., Tulsa, Oklahoma).

## Results

### Rapid, simultaneous evolution of dispersal propensity and ability

After 10 generations of selection, the VB populations were found to have significantly greater dispersal propensity (Fig. 1a, F_1,3_ = 148.1, P = 0.001) and dispersal ability (Fig. 1b, F_1, 3_ = 41.32, P = 0.008) compared to the VBCs. This suggests that compared to the controls, in the selected populations, a larger fraction of flies initiated dispersal and those dispersers travelled farther. It was interesting to note that only 10 generations of selection was sufficient to produce a significant divergence for these dispersal traits. We repeated the experiment after 20 generations of selection and the VB populations again had a significantly higher propensity (Fig. 1c, F_1,3_ = 22.68, P = 0.02) and ability (Fig. 1d, F_1,3_ = 68.8, P = 0.004) than the corresponding VBCs.

### Selected flies dispersed more even in the absence of stress

After 33 generations of selection, we again measured the dispersal traits of VBs and VBCs. The experimental set-up was identical to the previous assays mentioned above except each source now contained a supply of moisture and nutrition such that the flies were neither starved, nor desiccated. This removed two of the major proximate reasons for dispersal present during the process of selection. However, even in the presence of food in the source, the VB populations were found to have significantly greater dispersal propensity than VBCs (Fig. 1e, F_1,3_ = 60.78, P = 0.004) and ability (Fig. 1f, F_1, 3_ = 15.23, P = 0.03) compared to the VBCs.

Taken together, the above results imply that multiple components of dispersal had rapidly and simultaneously evolved in the selected populations, and this difference was observable irrespective of the presence or absence of a proximate reason for them to disperse. We next investigated the implications of these changes in dispersal components, on the spatial distribution of the organisms, i.e. the dispersal kernel (Nathan, et al. 2012).

### Evolution of dispersal kernel and increased frequency of LDDs in the selected populations

There was a clear difference between the distributions of the dispersal distances of the VBs and the VBCs (Fig. 2a), suggesting that the VB kernel had evolved due to selection. All the higher percentiles (75 onwards) of the dispersal kernel of VBs were greater than the corresponding percentiles of VBCs (Fig. 2b) which indicates the presence of a greater number of Long-Distance-Dispersers (LDDs) in the selected populations. This also suggested that the overall kernel shape has changed, which was supported by the fact that VB populations had a significantly greater standard deviation (Fig. 2c, MWU = 0.0, P = 0.02, *d* = 4.45), lesser positive skew (Fig. 2d, MWU = 0.0, P = 0.02, *d* = 1.79) and more negative kurtosis (Fig. 2e, MWU = 0.0, P = 0.02, *d* = 2.23) compared to the VBCs. For all these shape statistics, effect sizes of the differences between VB and VBC populations were large (i.e. *d*>0.8). When we fit a negative exponential distribution, (*y* = *ae^−bx^*) to the data (Fig. S3), we found that the values of the intercept parameter *a* (Fig. 3a, MWU = 0.0, P = 0.02, *d* = 3.77) and the slope parameter *b* (Fig. 3b, MWU = 0.0, P = 0.02, *d* = 4.17) for the VB kernels were significantly lower than the VBCs (see Table S1 for R^2^ values). This indicates a general flattening of the shape and fattening of the tail of the kernel in the selected populations.

**Figure 2.**
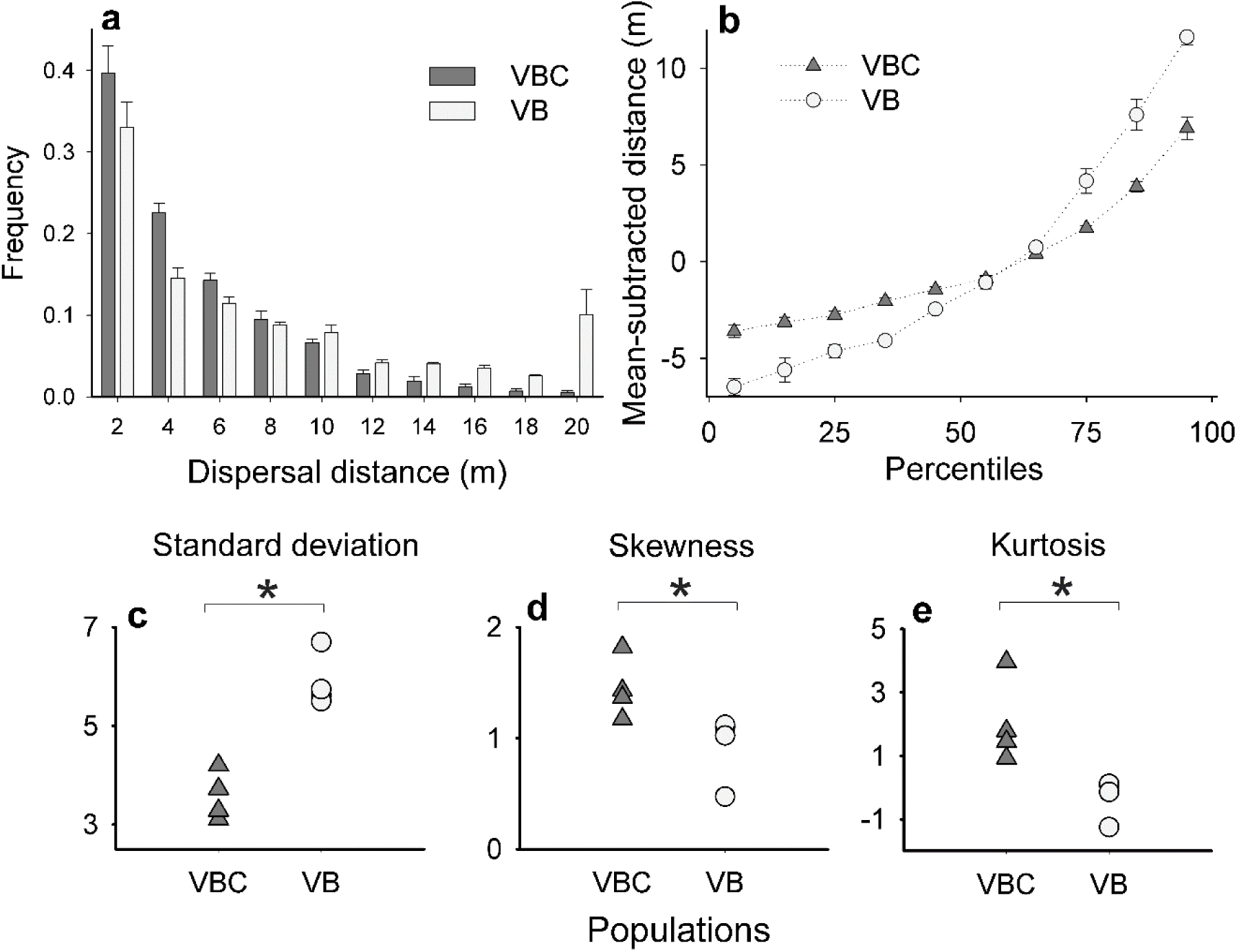
Evolution of location and shape parameters of dispersal kernel. (**a**) Overall dispersal kernel of VB and VBC populations. Each bar represents the average frequency of flies counted in the corresponding distance-bin for the four populations of each of VB and VBC. Error bars = standard error. (**b**) 5^th^ to 95^th^ percentile for the mean-subtracted kernels of VB and VBC populations. The error bars represent standard errors around the mean. In few cases the error bars are too small to be visible. Each percentile point represents data pooled over four VB or VBC populations each of which had three independent measurement replicates. (**c**) Standard deviation, (**d**) Skew and (**e**) Kurtosis of the dispersal kernels. Each point (triangle for VBC and circle for VB) represents data from one replicate population, pooled over three independent kernel measurements. Together these panels indicate that the dispersal kernels of VBs have become flatter and their tails have become fatter. * denotes P<0.05 for Mann-Whitney U-tests.

**Figure 3.**
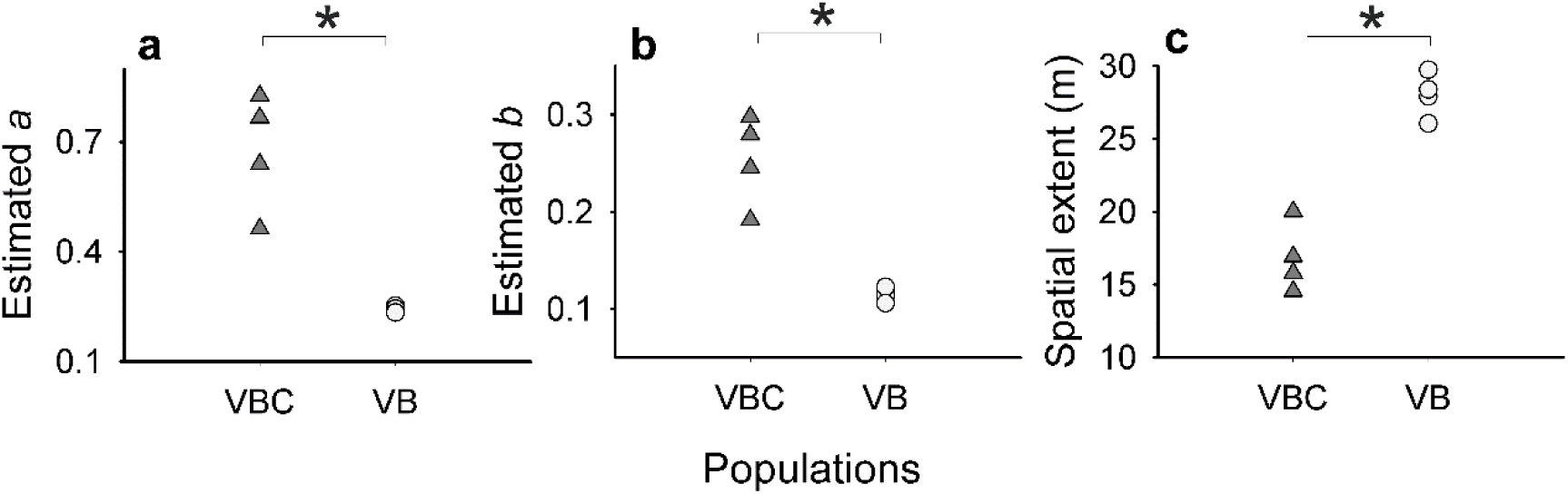
Evolution of the parameters of dispersal kernel and the spatial extent. Dispersal kernel parameters were estimated by fitting the negative exponential *y* = *ae^−bx^*, where *x* is the distance from the source and *y* is the frequency of individuals found at *x*. Estimated values of **(a)** *a* and (**b**) *b* are significantly lower for VBs than VBCs. (**c**) Using the fitted curve, spatial extent of each of VB and VBC populations was computed by finding the distance from the source, beyond which 1% of the population is expected to reach. Spatial extents of VBs were greater than VBCs, indicating an increase in LDDs in the population. Each point (triangle for VBC and circle for VB) represents data from one replicate population, pooled over three independent kernel measurements. * denotes *P*<0.05 for Mann-Whitney U-tests.

The mean spatial extent of the VBs and VBCs were found to be 28 m and 16.8 m respectively, which implies an increase of 67% (Fig. 3c, MWU = 0.0, P = 0.02; *d* = 5.67). In other words, if we compare the top 1% of the dispersing individuals, VBs travel ∼67% greater distance than VBCs. This implies an increase in the proportion of LDDs in the population (i.e. the fatness of the tail of the distribution). The fact that this increase was obtained after only 33 generations of selection suggests that evolvability of the kernel cannot be ignored in medium to long-range forecasts of phenomena like the rate of range expansion, disease spread and invasion speed.

It should be noted here that the Mann-Whitney U test (MWUT) compares the ranks of the observations across two groups (Zar 1999). This implies that for any number of tests, as long as the sample sizes and the relative ranks are the same, the U- and P-values will be identical. This is what is happening for the 6 MWUTs in Figs 2c-2e and Fig 3. There are absolutely no overlaps between the VB and VBC values in any of these figures, as a result of which, in all the MWUTs, the ranks for one group is 1,2,3,4 and that for the other is 5, 6, 7, 8. Not surprisingly, all of them yield exactly the same values of U and P.

In short, our results indicate that even if we account for the increased mean, the shape of the dispersal kernels of the VBs had evolved to be substantially different from that of the VBCs.

### *Drosophila* dispersal is sex-biased but both sexes respond similarly to selection

In presence of stress (performed after 20 generations of selection), males had greater ability to disperse than females (Fig. 4b, F_1,3_ = 16.28, P = 0.027), although in terms of dispersal propensity both the sexes performed equally (Fig. 4a, F_1,3_ = 3.47, P = 0.16). Interestingly, when assayed in the absence of stress after 33 generations, the trend reversed. Here we found that females had significantly lower propensity than males (Fig. 4c, F_1,3_ = 21.59, P = 0.019) but the ability of both the sexes was not different from each other (Fig. 4d, F_1,3_ = 2.23, P = 0.23). Given the presence of sex-biased dispersal in ability and propensity (albeit in different environments) we continued to investigate whether individuals of both the sexes responded equally to selection for dispersal. But we did not find any significant treatment × sex interaction in presence of stress with respect to dispersal propensity (Fig. S3a, F_1,3_ = 1.98, P = 0.25) or ability (Fig. S3b, F_1,3_ = 0.52, P = 0.52). In the absence of stress too, the interaction term was not significant in case of either propensity (Fig. S3c, F_1,3_ = 0.21, P = 0.68) or ability (Fig. S3d, F_1,3_ = 2.2, P = 0.24).

**Figure 4.**
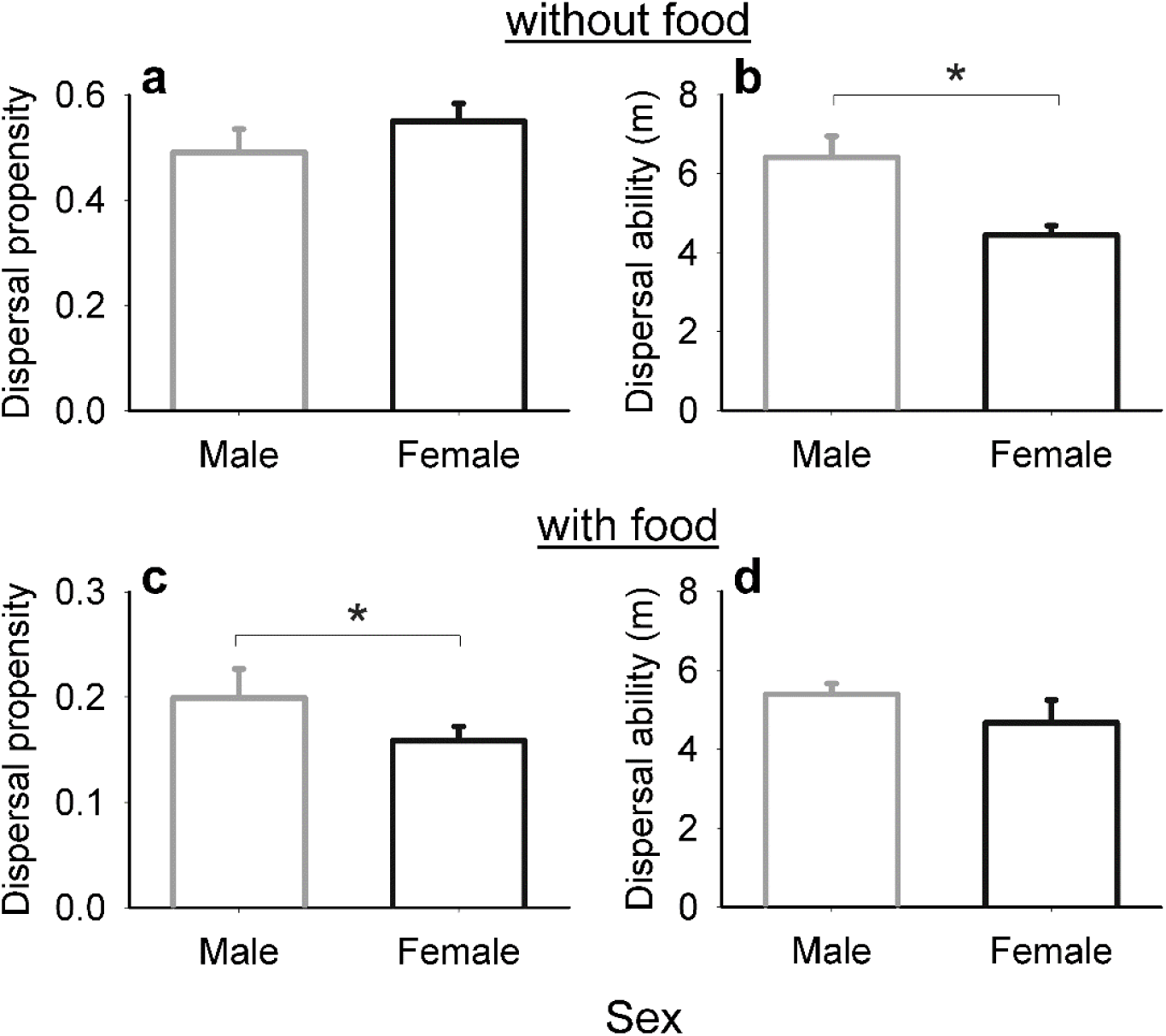
Sex-biased dispersal in the presence and absence of food. Dispersal (a) Propensity and (b) Ability in the absence of food in the source container. Dispersal (c) Propensity and (d) Ability in the presence of food in the source container. Males had significantly greater dispersal ability in the absence of food, but significantly higher propensity in the presence of food in the source. This shows that the expression of sex-biased dispersal can vary depending on which component is being measured and the environmental condition under which dispersal takes place. Error bars represent standard errors around the mean (SEM) and * denotes *P*<0.05 for the main effect of sex in the ANOVA.

## Discussion

During dispersal, various physiological and behavioral attributes pertaining to the different dispersal stages (i.e. emigration, travel and arrival/settlement) (Bowler and Benton 2005, Cote, et al. 2010) interact with each other and the external environment. These interactions can lead to a variety of costs for the organism (Bonte, et al. 2012). In order to evolve greater dispersal, organisms need to optimize over various physiological or genetic constraints to minimize the overall fitness cost of dispersal. For example, when the conditions are unfavorable and the cost of dispersal is less than the cost of staying back in the natal patch, enhanced dispersal propensity is likely to evolve (Friedenberg 2003). However, when the cost of travelling or settlement is very high, dispersal propensity might fail to evolve, even if there is reduced fitness in the natal patch (Cheptou, et al. 2008). As the number of factors and interactions that affect dispersal is very high, evolution of dispersal turns out to be a complex phenomenon. Thus, it becomes difficult to predict which components of dispersal would evolve and which would not, under the influence of various ecological circumstances.

Not surprisingly therefore, there is a large variation in the outcomes of experimental evolutionary studies on dispersal. For example, in spider-mites (*Tetranychus urticae*), it has been shown that dispersal propensity evolves when direct selection is imposed on dispersal rate (i.e. those who disperse early are selected) (Yano and Takafuji 2002). Interestingly, in the same model system, propensity fails to evolve when selection is imposed directly on propensity (Tien, et al. 2011) or on dispersal ability (Bitume, et al. 2011). Another study with the same model organism (Fronhofer, et al. 2014) showed that spatially-correlated extinctions favored the evolution of long-distance dispersers (LDDs) which is related to increased dispersal ability. However, in the same experiment, dispersal propensity did not evolve. This was because, positive spatial-correlation of extinction, in the absence of a significant increase in dispersal ability, substantially increases the cost of leaving the current habitat. A theoretical study on the evolution of passive dispersal of seeds on a fragmented landscape also suggests that spatial autocorrelation of nearby habitats can lead to the evolution of long-distance dispersal, but not propensity (Hovestadt, et al. 2001). To summarize, all these selection studies suggest that multiple components of dispersal (here propensity and ability) cannot evolve together.

### Multiple dispersal components can evolve simultaneously under condition-dependent selection

In our study, only the first 50% of the adults that reached the destination were allowed to breed. Thus, there was a direct selection for dispersal propensity (i.e. the tendency to leave the source patch). Moreover, as the length of the path increased over generations, there was also a direct selection on dispersal ability (i.e. the ability to travel the required distance). Consequently, within 10 generations of selection, dispersal propensity (Fig. 1a) and ability (Fig. 1b) of the selected lines (VBs) was significantly greater than the controls (VBCs). We measured these two components of dispersal again after 20 generations of selection and reached an identical conclusion (Figs 1c and 1d). Both these assays were performed under conditions similar to the selection (i.e. no food in the source) which increases dispersal propensity of the flies (*cf* Fig. 1c and Fig. 1e), presumably due to starvation and desiccation stress. Thus, the dispersal, in this case was *condition-dependent* (*sensu* Denno and Roderick 1992, Matthysen 2005), i.e. primarily driven by external cues.

The simultaneous evolution of dispersal propensity and ability in VBs was interesting because, in earlier studies, multiple components of dispersal had failed to evolve together (Bitume, et al. 2011, Fronhofer, et al. 2014, Tien, et al. 2011, Yano and Takafuji 2002). Our results also differ from theoretical (North, et al. 2011) and field (Baguette, et al. 2003, Cheptou, et al. 2008, Schtickzelle, et al. 2006) studies which predict that increased habitat fragmentation should have a negative effect on dispersal propensity. This apparent discrepancy is resolved when we observe that in some of these studies, the mortality during the travelling phase is so high that individuals with lower dispersal propensity have greater fitness even with habitat destruction (Baguette, et al. 2003, Cheptou, et al. 2008; although see Schtickzelle, et al. 2006). In our study, since ∼50% of the flies were able to reach the destination, the cost of dispersal was not prohibitively high. This allowed dispersal propensity to evolve, as predicted in some of the earlier theoretical studies (Heino and Hanski 2001, Zheng, et al. 2009).

### Phenotype-dependent dispersal can evolve even under condition-dependent selection

After demonstrating the evolution of condition-dependent dispersal, we next investigated the evolution of phenotype-dependent dispersal. This refers to dispersal tendencies that are intrinsic to the organisms (Clobert, et al. 2009), and thus independent of the dispersal cues. For this set of assays, we placed a food plate (which also provided moisture) in the source. Thus, the flies in the source experienced no starvation or desiccation stress, which removed the two major proximate reasons for dispersal experienced during the process of selection. Even under these conditions, a significantly greater proportion of VBs dispersed (Fig. 1e) to a larger average distance (Fig. 1f) than the VBCs.

This result is important because all previous experimental evolution studies have both imposed and assayed for phenotype-dependent selection (Bitume, et al. 2011, Fronhofer, et al. 2014, Keil, et al. 2001, Ogden 1970, Tien, et al. 2011, Yano and Takafuji 2002). The evolution of phenotype-dependent dispersal in such studies is intuitive. However, our results suggest that phenotype-dependent dispersal can evolve rapidly as a result of selection for condition-dependent dispersal. The presence of such constitutive dispersers will evidently affect the dispersal-related properties of a population which in turn can affect a large number of community- and ecosystem-level processes including range expansion (Travis and Dytham 2002), invasion (Kot, et al. 1996, Shaw and Kokko 2015), spread of diseases (Rappole, et al. 2006) and community dynamics (Leibold, et al. 2004). The other aspect of dispersal that would affect these processes is the dispersal kernel.

### Evolution of dispersal kernel and LDDs in the selected populations

In principle, two populations can have very different dispersal kernels because of underlying evolved differences in the phenotype, or the environment, or both. Thus, in order to demonstrate the evolution of the kernel due to phenotypic evolution in the populations, it is important to control for the environment in which dispersal is assayed. We achieved this in our study by making the path through which the flies moved completely homogeneous and devoid of any potential environmental feature (like food or predators). More importantly, both the VBs and VBCs experienced the same environment during dispersal. Thus, all the differences observed between the kernels of these two populations were attributable to the underlying phenotypic differences. Evidently, we cannot claim this kernel to be the “natural kernel” of the flies as there are potentially infinite numbers of such “natural kernels” (one for every environmental state). However, this study showed that for a given environment, condition-dependent selection for dispersal can alter the location and shape of the kernel (Fig. 2 and Figs. 3a-3b) and enhance the fraction of LDDs in the population (Fig. 3c), even when there is no proximate reason for dispersal. This is an interesting point because the shape of the dispersal kernel is often considered to be a static entity in much of the theoretical literature (Chapman, et al. 2007, Krkošek, et al. 2007). Our results are thus in line with more recent theoretical (Phillips, et al. 2008, Starrfelt and Kokko 2010) and empirical studies (Fronhofer, et al. 2014) that considered the possibility of evolving kernel shapes.

Our results also showed that the skew and kurtosis of the selected populations had reduced compared to the controls (Fig. 2d and Fig. 2e) which is consistent with observations on invasive cane toad populations in Australia and results of mathematical modelling on the same species (Phillips, et al. 2008). For both studies, the change in the kernel shape parameters can be attributed to the increased frequency of LDDs in the population.

### Sex-biased dispersal exists in *Drosophila*

Although there are many examples of sex-biased dispersal (SBD) among birds and mammals (reviewed in Pusey 1987), relatively few cases of SBD have been reported among insects like butterflies (Bennett, et al. 2013) and ground beetles (Lagisz, et al. 2010). It has been shown that *Drosophila pachea* exhibits SBD while *D*. *nigrospiracula* and *D. mojavensis* do not (Markow and Castrezana 2000). However, to the best of our knowledge, no prior study has looked at SBD in *Drosophila melanogaster*. Therefore, our first aim was to see whether SBD exists at all in this species.

When there was no food in the source, the males had a greater ability to disperse (Fig. 4b), but there was no sex-bias in dispersal propensity (Fig. 4a). Interestingly, the situation was reversed when there was food in the source, i.e. there was no difference in ability (Fig. 4d), but the males had greater dispersal propensity (Fig. 4c). These results highlight two major issues in studying SBD. First, the presence of SBD for any one component of dispersal is no guaranty for the presence of SBD for another dispersal component (Clutton-Brock and Lukas 2012). Second, the fact that SBD for propensity and ability were seen in the absence and presence of food respectively, illustrates that the manifestation of SBD for a given dispersal component can be condition-dependent. Taken together, these observations suggest that across-study comparisons of SBD are not possible, until and unless they refer to the same dispersal component, under similar environmental conditions.

One potential complication with these experiments is that the dispersal assays in presence and absence of food were not conducted at the same time. Thus, in principle, it is possible that the differences between the results in presence and absence of food are due to the selection that happened during the intervening time. Although we failed to come up with any biological reasoning, we could not logically rule it out either. Note that our first observation (SBD for one component does not guaranty SBD for another) remains unaffected by this complication.

### Both sexes respond similarly to selection for dispersal in *Drosophila*

Prior studies have shown that selection can lead to sex-specific effects on dispersal-related traits (Legrand, et al. 2016). Therefore, our next major question was whether selection had made dispersal more sex-biased in the VB populations. This kind of a bias was expected because of an inherent asymmetry in our selection protocol: non-dispersing males could, in principle, pass their genes to the next generation by impregnating dispersing females, while the females had no such option in terms of their evolutionary contribution. This implied a potentially stronger selection pressure on the females for dispersal-related traits, which could lead to a sex × selection interaction for propensity or ability. In short, VB females were expected to diverge more from the VBC females than the VB males from the VBC males.

The sex × selection interaction was not statistically significant irrespective of the presence or absence of food (Fig. S3). Unfortunately, we could not conduct post-hoc tests for these differences, as the sex × selection effect was not significant in either ANOVA. However, it can be safely said that there was no evidence to support that the males had responded less to selection for either dispersal ability or propensity. There can be at least two potential (and mutually non-exclusive) reasons for this observation. First, in *Drosophila*, there is substantial evidence for last male precedence, i.e. when the females mate multiple times, the last male to mate sires more offspring (reviewed in Parker 1970). Thus, the males that mated with the females after dispersal could have had much greater fitness than those that mated before dispersal. This could considerably increase the selection pressure on the males to disperse and explain why the males maintained their advantage in terms of propensity and ability in the VB populations. The other possibility is that dispersal traits are controlled by the same loci in both females and males in *D. melanogaster*, such that it is not possible for the sexes to respond differently to selection for dispersal. Interestingly, in terms of trends, the VB males always had greater dispersal propensity and ability than the VBC males (Fig. S3). This suggests that, irrespective of its relative magnitude with respect to the females, there was substantial positive selection pressure on VB males for dispersal components.

### Caveats

In this study, we selected for ambulatory dispersal in fruit flies. It is well known that this is not the primary mechanism by which fruit flies disperse in nature (Dobzhansky 1973) and we had no intention of examining that topic in a laboratory study. Our aim here was to investigate, given a mode of movement, how various aspects of dispersal interact and evolve. One of the reviewers have pointed out that in our kernel assays the flies had not attained their equilibrium distribution of dispersal distances after six hours, which could potentially invalidate our kernel measures. We respond to this criticism in some detail in Online Appendix S1: Text S2 and show why this concern is misplaced.

### Implications of our results

There is a growing realization that multiple components must be investigated simultaneously to obtain a complete picture of dispersal evolution (Bonte, et al. 2012). However, there is no theoretical or empirical expectation about the relationship between the various dispersal components, i.e. evolution of propensity does not let us predict anything about the evolution of ability and vice versa. Given this scenario, our result about the concurrent evolution of multiple dispersal components can be taken as a null model. In other words, whenever a particular component of dispersal is seen not to evolve, elucidating the reasons for that can become the focus of an investigation. Furthermore, our study shows that under gradual directional selection of moderate intensity and in the absence of conflicting selection pressures, dispersal can evolve rapidly, and substantially. Such conditions are expected to be fairly common in nature, particularly in regions where climate changes or habitat degradations are gradual but steady. More critically, our results indicate that once evolved, these traits can express themselves even in absence of the proximal stresses (i.e. become phenotype-dependent). This could lead to organisms with intrinsically high rates of dispersal. On one hand, this could reduce the chances of local extinction (Brown and Kodric-Brown 1977, Forney and Gilpin 1989) and ensure greater gene flow among populations (Vilà, et al. 2003). While on the other, this could increase the invasiveness of species (Kot, et al. 1996, Neubert and Caswell 2000), increase the rate of spread of diseases (Keeling, et al. 2001) and induce instability in metapopulation dynamics through enhanced synchrony between neighboring subpopulations (Dey and Joshi 2006). Figuring out the magnitude by which different dispersal components evolve and how that affect these ecological processes will be a major challenge not only for ecologists but also for ecological economists and conservation biologists (Buoro and Carlson 2014).

## Acknowledgements

We thank Adithya E Rajagopalan for help in running the experiments, Selveshwari S. for help in preparing the figures and Emanuel Fronhofer for insightful comments.ST and AM were supported by Senior and Junior Research Fellowships respectively of the Council for Scientific and Industrial Research, Government of India. PMS and MAS were supported through the INSPIRE fellowship of the Department of Science and Technology, Government of India. VRSS was supported through the GE Foundation Scholar Leaders Program. This study was supported by a research grant (#EMR/2014/000476) from Department of Science and Technology, Government of India and internal funding from IISER-Pune. The authors declare no conflict of interest.

## Online Appendix S1

### Text S1 Supplemental Experimental Procedures

#### 1 Ancestral populations

The experimental populations used in this study were derived from four independent large (breeding size of ∼2400) laboratory populations of *Drosophila melanogaster* (DB_1-4_) which in turn trace their ancestry to four outbred populations called JB_1-4_. The detailed maintenance regime and ancestry of the JB1-4 populations has been described elsewhere (Sheeba, et al. 1998). The maintenance regime of the DB_1-4_ populations are similar to the JB_1-4_, except that the former set of flies are introduced into population cages on the 12^th^ day after egg collection

From each DB_*i*_ population (where *i*∈[1, 4]), we derived two populations: VB_*i*_ (short for ‘vagabond’, subjected to selection for dispersal) and VBC_*i*_ (corresponding no-dispersal control). Thus VB and VBC populations that share a numerical subscript (e.g. say VB_1_ and VBC_1_) were related by ancestry (DB_1_ in this case), and hence were always assayed together and treated as blocks in statistical analyses.

#### 2 Maintenance regime of experimental populations

The adults of both VBs and VBCs were maintained in plexi-glass population cages (25 cm × 20 cm × 15 cm) at a high adult number (∼2400 individuals) to avoid inbreeding. Following earlier protocols, both the larvae and the adults were maintained at 25°C and constant light conditions (Sheeba, et al. 1998). The flies were made to oviposit on petri-plates containing banana-jaggery medium for 12-16 hours. After oviposition, we cut small strips of the medium, each containing ∼60–80 eggs, and introduced them individually into 35 ml plastic vials that had ∼6 ml of the same banana-jaggery medium. This ensured that the larvae were raised under low to moderate level of crowding, and there was no confounding effect of density-dependent selection (Joshi 1997). The adults started emerging by the 8^th^ – 9^th^ day after egg collection and on the 12^th^ day, the VB populations underwent selection for dispersal (see below). Since at 25°C temperature, all normally developing adults eclose by 10^th^ – 11^th^ day, our selection protocol ensured that there was no inadvertent selection for faster larval development (Prasad, et al. 2001). After the imposition of selection, the flies were transferred to the population cages and immediately supplied with excess live yeast-paste to boost their fecundity. Around 40 hours after this, the flies were supplied with a fresh petri-plate containing banana-jaggery medium for oviposition. The eggs so collected formed the next generation and the egg-laying adults were discarded, ensuring that adults from two different generations never co-exist. Thus, both VBs and VBCs were maintained under 15-day discrete generation cycles. For each VB population, we collected eggs in 80 vials (thus leading to approximately 4800 adults) while for VBCs, the corresponding number of vials was 40. This ensured that after selection (see next section), the breeding population of the VB populations was similar to that of the VBCs.

#### 3 Selection protocol

The apparatus for selection for dispersal consisted of three components: a *source*, a *path* and a *destination*. The source was an empty transparent cylindrical plastic container of diameter 11 cm and height 16 cm with a funnel attached to one end (Fig. S1). The diameter of the broad end of the funnel matched that of the source, while the diameter of the exit to the stem was 1.8 cm. The path connecting the source with the destination consisted of a transparent plastic tube of inner diameter ∼1 cm. The destination too was a cylindrical plastic container (diameter 11 cm and height 16 cm) and contained a supply of moisture in the form of a strip of wet cotton. The end of the path protruded ∼10 cm inside the destination (Fig. S1). This protrusion helped in reducing the rate of backflow as, after getting out of the path, the flies typically spent most of their time on the walls or floors of the container, and hence mostly failed to locate this aperture. To make the overall setup compact, the path was coiled (in the horizontal plane). The length of the path was 2 m at the beginning of the selection, but was increased intermittently. By generation 33, when the last set of assays were performed, the path length had reached 10 m.

In order to impose the selection, on the 12^th^ day after egg-collection, ∼2400 adults (coming out of 40 vials) of a given VB_*i*_ population were placed in a source, which was then connected to the destination via the path. The entire setup was placed in a well-lit room maintained at 25 °C. Since the source had no moisture, the flies experienced desiccation. Pilot runs with the ancestral DB populations had shown that under these environmental conditions, a subset of the flies tended to move through the opening towards the destination. Pilot studies also showed that very few flies dispersed in the presence of food in the source and therefore we decided to impose selection in the absence of food. The flies were allowed to disperse for six hours or until roughly 50% of the population reached the destination (whichever happened earlier). The arbitrary cut-off of six hours was chosen because assays in the lab had demonstrated that under desiccating conditions, there was almost no mortality during the first six hours (S. Tung personal observations). Only the flies that reached the destination were allowed to breed for the next generation. Since the imposed selection allowed ∼50% of the flies to breed, there were two independent “source-path-destination” setups, with ∼2400 flies in the source, for each VB_*i*_ population. Post-selection, the dispersed flies in the two destination containers for a given VB_i_ population were mixed and transferred to a population cage. They were then supplied with live-yeast paste and after ∼40 hours, eggs were collected (as mentioned above). The VBCs were maintained similarly as the VBs except two major differences. Firstly, after transferring the flies into the source, the exit was blocked by a cotton plug and the flies were allowed to desiccate for 3 hours or till 25% of the VBs reached their destination (whichever was earlier). Following the protocol for the VB flies, the VBC flies were then supplied with a moist cotton plug for the remaining duration of VB dispersal. This controlled for the inadvertent desiccation experienced by the VB flies in the source and the path, as part of the selection protocol. It should be noted here that there was almost zero mortality in the VBC flies during this time, thus ensuring that the selection pressure for desiccation resistance was at best, mild. Secondly, all the flies in the VBC populations were allowed to breed, thus ensuring no selection for dispersal.

#### 4 Assays

All assays were performed after relaxing the selection on both VB and VBC populations for one generation. For this, the VB and VBC flies were transferred directly into the corresponding cages on the 12^th^ day after egg collection. The progeny of these flies, henceforth referred to as the relaxed populations, were used for the assays. This common-rearing ensured that influence of phenotypic plasticity or non-genetic parental effects were ameliorated. Additionally, to remove any extraneous influence due to larval crowding, egg density was kept to ∼50 eggs on ∼6mL food in each vial. Furthermore, as the assays for each of the four blocks required us to sex and count ∼12,000 flies, it was not logistically possible to assay more than two blocks in a given generation. Therefore, each assay was conducted over two successive generations and it is the latter value which is reported in the paper (i.e. for the *t^th^* generation assay, VB_1_-VBC_1_ and VB_2_−VBC_2_ were assayed in generation t-1 while VB_3_-VBC_3_ and VB_4_-VBC_4_ were assayed in generation t). For example, for the 33rd generation assays, block 1 and 2 were assayed during the 32^nd^ generation of selection, while block 3 and 4 were assayed during the 33^rd^ generation and so on. This is not a problem in terms of our statistical analysis as block was explicitly recognized as a random factor in our ANOVA.

#### 4.1 Dispersal assay in presence and absence of food

This assay was performed thrice-after 10, 20 and 33 generations of selection in order to assess the difference in dispersal propensity and ability between the VBs and the VBCs. The assay-setup was similar to the selection setup (see ‘Selection protocol’ above and Fig. S1) except for the length of the path. The path-length was 10 m for the assay performed after 10 generations of selection and for the rest it was 20 m. Furthermore, to obtain the distribution of the location of the files after dispersal, the path was divided into multiple detachable sections: 20 sections of length 0.5 m each for the first 10 m and followed by 10 sections of length 1 m each for the rest (1 m sections were not present when the path-length was only 10 m). The destination container (a 250 ml plastic bottle) did not contain food or water but had a long protrusion of the path into it, to reduce the backflow of flies. On the 12^th^ day after egg collection, ∼2000 adult flies were introduced into the source container and were allowed to disperse for 6 hours. During this interval, the entire setup was kept undisturbed under constant light and at a temperature of 25˚C. After the end of dispersal run, the setup was dismantled; the openings of the source, the destination, and each section of the path were secured carefully with cotton plugs, and labelled appropriately. The flies were then heat killed and the location (in terms of the distance from the source) and sex of each fly was recorded. For each VB_*i*_ and VBC_*i*_ population, there were three such replicate kernel setups.

We performed two kinds of kernel assays: a) with an empty source and b) in the presence of ∼20 ml banana-jaggery medium in the source. The former set of assays was performed after 10 and 20 generations of selection while the latter set of assays happened after 33 generations of selection. In total, these assays involved segregating according to sex and scoring of ∼140, 000 flies.

### 5 Dispersal components

#### 5.1 Dispersal propensity

The proportion of total flies in the source that initiated dispersal was taken as the dispersal propensity (Friedenberg 2003). Thus propensity = (Number of flies found outside the source/ Total number of flies).

#### 5.2 Dispersal ability

The dispersal ability was computed only on the flies that left the source, based on the section of the path in which they were found after 6 hours. All flies found in a given section of the path were deemed to have travelled the distance between the source and the midpoint of the section. The destination container was considered as a part of the last path-section. Thus mathematically,

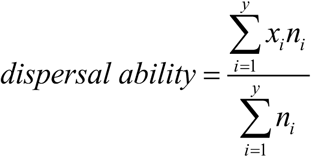

where, *n*_*i*_ is the number of flies found in the *i*^th^ path-section, *x*_*i*_ is the distance of the mid-point of this section from source and y is the total number of path-sections (here *y* = 30, see Section ‘Dispersal kernel assay in presence and absence of food’ for details). Since dispersal ability is measured only on the flies that came out of the source, the measure of propensity and ability were independent of each other.

### 6 Curve-fitting for estimating population extent

The data obtained from the dispersal kernel assay in presence of food, were fitted with the negative exponential distribution *y* = *ae^−bx^*, where *x* is the distance from the source, *y* is the frequency of individuals found at *x*, and *a, b* are the intercept and slope parameters respectively. For this we pooled the data of the three replicates for each of the four populations of VB and VBC, estimated the frequency for each distance, natural log-transformed all values and fitted the equation ln(y) = ln(a) – bx using linear regression. The estimated R^2^ values (Supplementary Table S1) ranged between 0.67 and 0.99 and the residuals showed no major trends. The value of population extent was estimated as *b*^−1^. ln (*a*/0.01), i.e. the distance from the source beyond which 1% of the population is expected to disperse.

During the linear regression, we observed that one data point in the kernel of the VB_3_ population seemed to be an outlier. Excluding this point from the kernel considerably improved the fit (R^2^ = 0.26 became R^2^ = 0.91) and the distribution of the residuals improved considerably. However, removing this outlier reduced the mean value of the spatial extent of VBs from 32.6 m to 28.01m. Incidentally, there were no changes in terms of the statistical significance in the Mann-Whitney U-tests for *a*, *b* or the spatial extent irrespective of whether the outlier is included or excluded. Therefore, in this study, we chose to report the value of population extent omitting the outlier. Note that this removal only makes our estimate of the spatial extent of VBs more conservative.

## Text S2 Estimation of dispersal kernel in this study

Dispersal kernel has been defined as the "distribution of the post-dispersal locations relative to the source point" (Nathan, et al. 2012). However, for constantly moving organisms like our fruit flies, it is impossible to define when dispersal has ended, particularly when they are still in the path. That is why we had to impose a temporal cut-off for the kernel assays, which is consistent with similar empirical studies in the literature (Bitume, et al. 2011, Markow and Castrezana 2000, Ogden 1970). As a reviewer has pointed out, since the flies were still active after six hours, it is possible that the measured distribution had still not reached equilibrium and therefore the kernel can be “linked to dispersal speed or the time needed to leave a patch, and not to dispersal distance per se”. This problem is intrinsic to the study of the dispersal kernel of any actively moving organism that does not settle to make nests or occupy territories, and there is no way to obtain the equivalent of an equilibrium “distribution of the post-dispersal locations” for such animals. Moreover, we still believe that the kernel that we measured gives valuable information about dispersal evolution, for the following reasons. First, our study compared the dispersal of the VBs and the VBCs under identical conditions which means that all comparative statements about the various aspects of the kernel (Figs 2 and 3) are valid, irrespective of whether the kernels were static or not. Second, looking at the individual kernels (Fig S2) it is clear that there is relatively little variation across the three replicates for any given population (VB or VBC). The same is true for the various shape parameters across the four replicates of the VBs or the VBCs (Figs 2c-e and 3). This suggests that even if the populations have not attained their equilibrium distribution of dispersal distance, they are probably fairly close to it. This is not surprising as other experiments in our laboratory have shown that ∼90% of the flies that leave the source in VBs and VBCs, do so within the first 90 minutes (Tung et al, manuscript under preparation). This implies that most of the flies spend ≥4.5 hours on the path, which is ∼66% of the total dispersal time. To summarize, we believe that even if we cannot demonstrate that we have measured dispersal kernel at equilibrium, this is a problem inherent with most active dispersers, it does not change any of the conclusions of our study, and our measured kernels are probably very close to the equilibrium any way.

**Table S1.** R^2^ values of the fitted kernels

| **Populations** | **$R^2$** |
| --- | --- |
| VB1 | 0.67 |
| VB2 | 0.73 |
| VB3 | 0.91 |
| VB4 | 0.65 |
| VBC1 | 0.97 |
| VBC2 | 0.97 |
| VBC3 | 0.99 |
| VBC4 | 0.98 |

**Figure S1.**
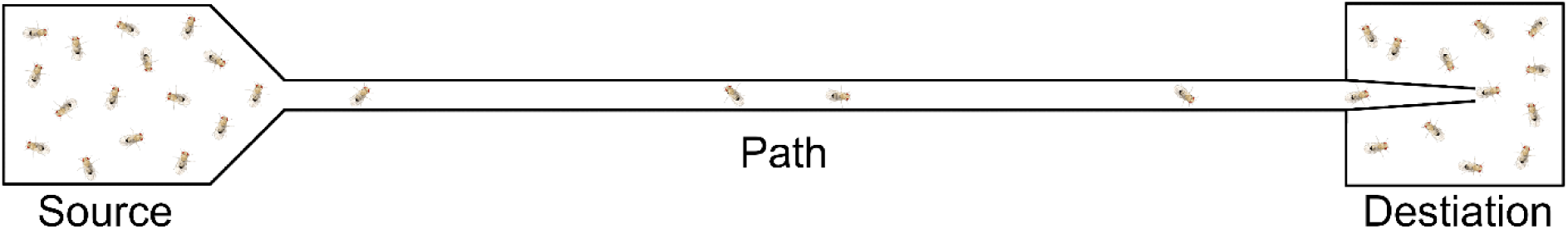
Schematic diagram of the selection and assay setup. The source and the destination are transparent plastic containers. The path is a transparent plastic tube. The path protrudes ∼10 cm inside the destination; this protrusion considerably reduces backflow of the flies. Here, all the three parts-- the source, path and the destination are detachable. The tiny objects oriented randomly inside the setup denote the flies. The length of the path increased from 2m to 10m during the 33 generations of selection reported here.

**Figure S2.**
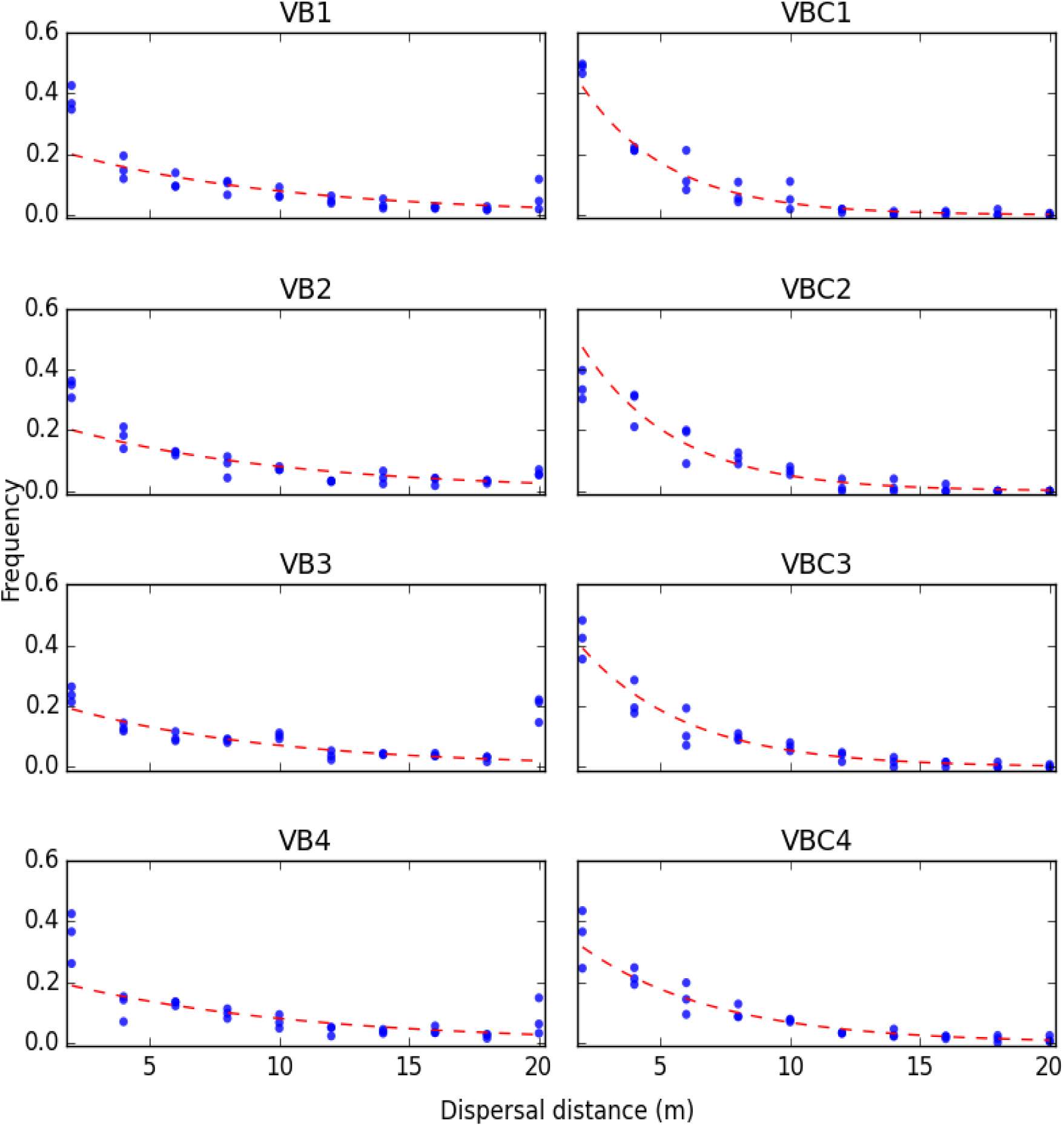
Fitted kernels of VB and VBC populations. In each panel, the frequency of individuals dispersed (scaled by total number of dispersed individuals) is plotted against the corresponding dispersal distances for each population. The three points at each dispersal distance correspond to the three measurement replicates of a population. The red dashed line is the negative exponential curve fitted to the pooled data of the corresponding population. Note, for VB3, the frequency value at dispersal distance 20 was considered as outliers and not considered during fitting.

**Figure S3.**
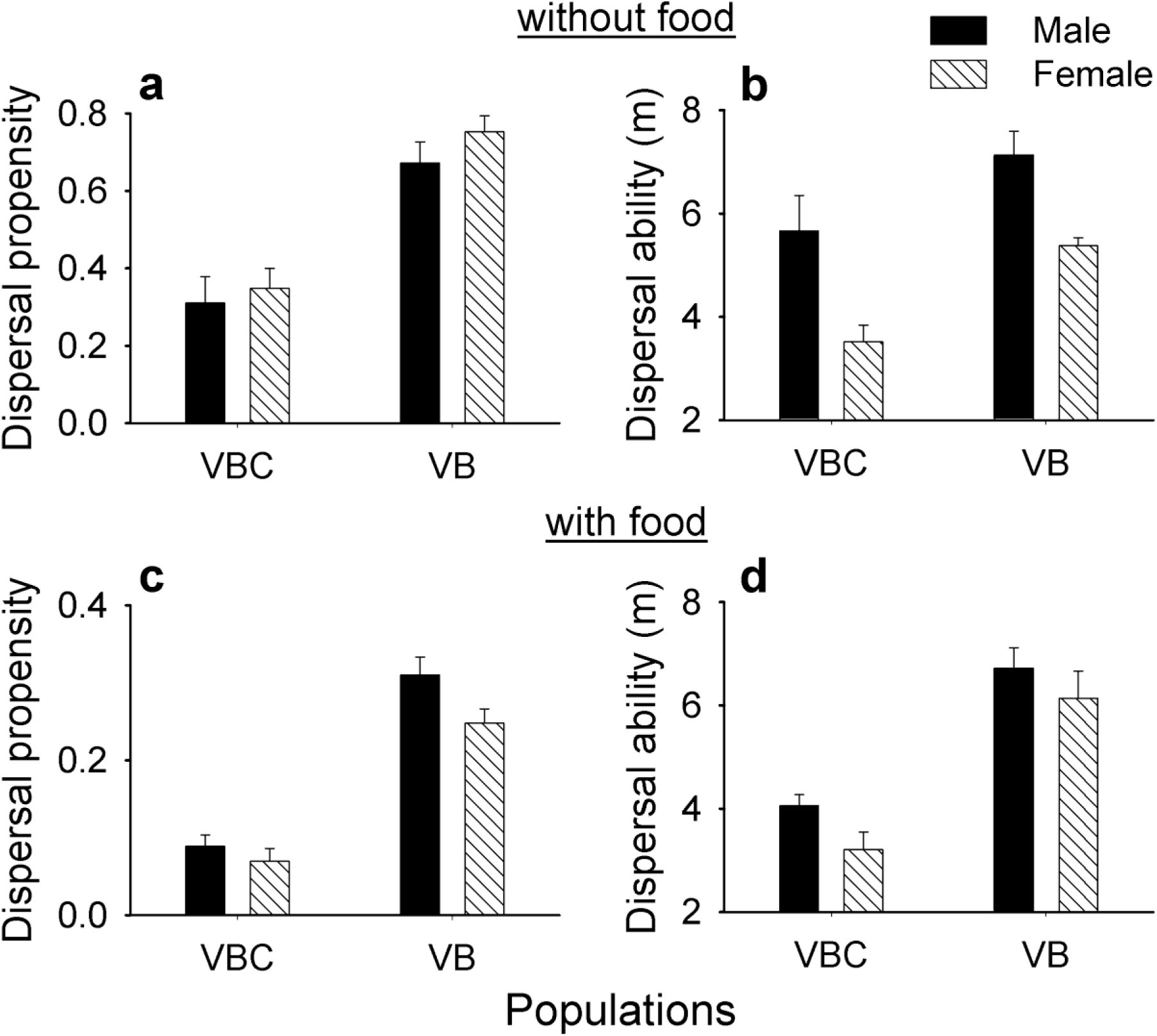
Selection × Sex interaction for dispersal propensity and ability in the presence and absence of food. Here none of the sex × selection interactions were statistically significant, thus we could not conduct post-hoc tests for pairwise comparisons of each groups. However, dispersal propensity (**a**) and ability (**b**) of both the sexes of VB populations were greater than those in VBCs in absence of food in the source. Similarly, even in presence of food in the source, males and females of VB populations had higher dispersal propensity (**c**) and ability (**d**) than VBCs. This showed that both sexes in VB responded equivalently to the selection for dispersal. The error bars represent standard errors around the mean (SEM).

